# VIRIDIC – a novel tool to calculate the intergenomic similarities of prokaryote-infecting viruses

**DOI:** 10.1101/2020.07.05.188268

**Authors:** Cristina Moraru, Arvind Varsani, Andrew M. Kropinski

## Abstract

Nucleotide based intergenomic similarities are useful to understand how viruses are related with each other and to classify them. Here we have developed VIRIDIC, which implements the traditional algorithm used by the International Committee on Taxonomy of Viruses (ICTV), Bacterial and Archaeal Viruses Subcommittee, to calculate virus intergenomic similarities. When compared with other software, VIRIDIC gave the best agreement with the traditional algorithm. Furthermore, it proved best at estimating the relatedness between more distantly related phages, relatedness that other tools can significantly overestimate. In addition to the intergenomic similarities, VIRIDIC also calculates three indicators of the alignment ability to capture the relatedness between viruses: the aligned fractions for each genome in a pair and the length ratio between the two genomes. The main output of VIRIDIC is a heatmap integrating the intergenomic similarity values with information regarding the genome lengths and the aligned genome fraction. VIRIDIC is available at viridic.icbm.de, both as a web-service and a stand-alone tool. It allows fast analysis of large phage genome datasets, especially in the stand-alone version, which can be run on the user’s own servers and can be integrated in bioinformatics pipelines. VIRIDIC was developed having viruses of *Bacteria* and *Archaea* in mind, however, it could potentially be used for eukaryotic viruses as well, as long as they are monopartite.

## Introduction

Intergenomic comparisons are useful in determining how viruses are related to each other. Indeed, the primary classification technique used by the International Committee on Taxonomy of Viruses (ICTV), Bacterial and Archaeal Viruses Subcommittee is based upon overall nucleic acid sequence identity. For a number of years a crude method of estimating this was derived from BLASTN searches at NCBI, by multiplying the “Query Cover” by the “Per. Ident” values. The Subcommittee established thresholds for the demarcation of viruses into species (95%) and into genera (~70%). While this technique is useful for undertaking pairwise comparisons, it is not convenient for comparisons of larger datasets.

There are a number of online tools and stand-alone software packages which have been used to compare viral genomes, including with the purpose of taxonomic classification. These include Average Nucleotide Identity (ANI; ANI Calculator; (Goris *et al.* 2007; Yoon *et al.* 2017; Han *et al.* 2016), OrthoANI (Lee *et al.* 2016), EMBOSS Stretcher (Ceyssens *et al.* 2009), Gegenees (Agren *et al.* 2012), JSpeciesWS (Richter *et al.* 2016), KI-S tool (https://f1000research.com/posters/7-147), PAirwise Sequence Comparison (PASC; (Bao *et al.* 2012; Bao *et al.* 2014), Sequence Demarcation Tool (SDT; (Muhire *et al.* 2014), Simka (https://arxiv.org/abs/1604.02412), Yet Another Similarity Searcher (YASS; (Noé and Kucherov 2005). Some of the tools not only calculate, but also offer a visual of the comparison of the nucleic acid sequence relatedness (reviewed in (Mahadevan 2016). To these we can add progressiveMauve (Darling *et al.* 2010) and VICTOR (Meier-Kolthoff and Göker 2017), which are focused on visualization of the alignments / genome relatedness, without explicitly giving access to the similarity values themselves.

The alignment algorithms used to calculate intergenomic relatedness vary from those based on the Needleman-Wunsch global alignment (Stretcher, SDT, PASC), to those based on BLASTN, either with previous genome fragmentation (Gegenees, OrthoANI) or without (PASC, VICTOR). With the exception of PASC and VICTOR, which can normalize the intergenomic identities to the whole genome length, most of the other tools normalize the intergenomic identities to the length of the alignment. This can lead to artificially high similarity values. The differences in algorithms can result in significant differences between the similarities reported by the different tools and can lead to inconsistencies in virus classification.

Here, with the purpose of offering a standardized and high-throughput tool for comparing viral genomes, we developed Virus Intergenomic Distance Calculator (VIRIDIC). VIRIDIC builds and improves on the traditional BLASTN method used by Bacterial and Archaeal Viruses Subcommittee from ICTV, to both calculate and visualize virus intergenomic relatedness. VIRIDIC is available as a web-service and as a stand-alone program for Linux, both accessible at viridic.icbm.de. It reports either intergenomic similarities or intergenomic distances.

## Methods

### VIRIDIC – development and workflow

VIRIDIC was developed in R 3.5 programming language (R Core Team 2018). The web interface was developed under the shiny web application framework (https://cran.r-project.org/web/packages/shiny/index.html). The stand-alone tool for Linux was wrapped in a container using the Singularity v. 3.5.2 software (https://sylabs.io/). This VIRIDIC version can be deployed on any systems running the Singularity software, without any additional installation and configuration steps.

VIRIDIC’s work flow consists of four steps. First, each viral genome is aligned against all other genomes in the dataset, using BLASTN 2.9.0+ from the BLAST+ package (Camacho *et al.* 2009) with the core parameters “-evalue 1 -max_target_seqs 10000 -num_threads 6”. The default alignment parameters are “-word_size 7 -reward 2 -penalty −3 –gapopen 5 –gapextend 2”. The user can choose between 3 other parameter sets: “-word_size 11 -reward 2 -penalty −3 –gapopen 5 –gapextend 2”, “-word_size 20 -reward 1 -penalty −2”, “-word_size 28 -reward 1 -penalty −2”.

Second, the BLASTN output is used to calculate pairwise intergenomic similarities. For one genome pair, the number of identical nucleotide matches reported by BLASTN is summed up for all aligned genomic regions. In the case of overlapping alignments, the overlapping part is removed from one of the aligned regions, such that, at the end, the different genome regions are represented only once in the alignments. The intergenomic similarity or distance is calculated as described below, as previously proposed (Meier-Kolthoff and Göker 2017).

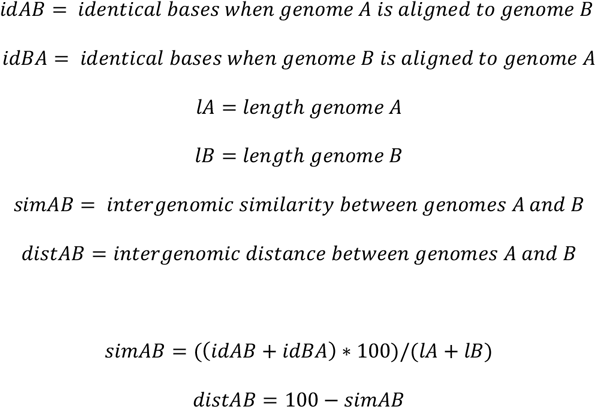

The intergenomic similarity algorithm has been implemented to run on multiple central processing unit cores using the future vs 1.17.0 R package (https://github.com/HenrikBengtsson/future).

In the second step, VIRIDC also calculates for each genome pair three other indicators related to the alignment: the aligned fraction for genome 1, the length ratio between genome 1 and genome 2, and the aligned fraction for genome 2.

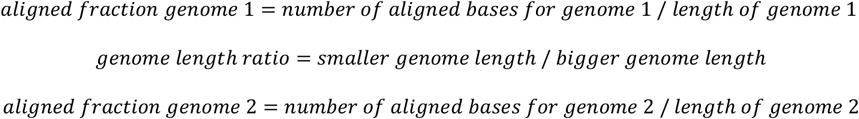

Third, VIRIDIC performs a hierarchical clustering of the intergenomic similarity values. For this, the intergenomic similarities are transformed into euclidian distances using the parallelDist v 0.2.4 R package (https://github.com/alexeckert/parallelDist), then clustered using the fastcluster v 1.1.25 R package (https://cran.r-project.org/web/packages/fastcluster/index.html) (Müllner 2013). For clustering, VIRIDIC uses by default the “complete” agglomeration method (see hclust function, fastcluster package). Several other agglomeration methods from the fastcluster package can be given as parameter.

Fourth, VIRIDIC graphically represents the intergenomic similarity values, the aligned ratios 1 and 2 and the genome length ratios as a heatmap, using the ComplexHeatmap v. 2.5.3 R package (Gu *et al.* 2016). The heatmap is ordered based on the genome clustering by their similarity values.

VIRIDIC outputs an ordered similarity / distance matrix (tab separated text format), a heatmap (pdf format) and a table with the viral genomes tentatively clustered at the species or genus level (tab separated text format). Additionally, the stand-alone tool offers access to several intermediary files, both in RDS and tab separated text format, containing further information about the alignments. These files could be eventually be integrated in bioinformatics pipelines.

### Benchmarking VIRIDIC

The dataset used for benchmarking consisted of 60 T7-like phages genomes, from the *Autographviridae* family, downloaded from the GenBank RefSeq database (Haft *et al.* 2018). These genomes were chosen because they are related, colinear and have an average genome size of 39.4 kbp (range: 31.5 – 41.7) and G+C mol% content of 50.7 (range: 42.6 – 61.8; SI Table 1). The testing dataset also contained the Pelagibacter phage HTVC011P genome, used as outlier for the T7-like phages. For this dataset of 61 phages, the intergenomic similarities where calculated with the following tools: Sequence Demarcation Tool (SDT) (Muhire *et al.* 2014), Pairwise Sequence Comparison (PASC) (Bao *et al.* 2012), OrthoANI (Lee *et al.* 2016), Gegenees (Agren *et al.* 2012), and VIRIDIC.,

Additionally, two *Salmonella* phages (GE_vB_N5 and FE_vB_N8) were used for the illustration of genome and alignment length differences. The genome of the K155 strain of the T7 phage was used to test the effect of genome permutations and reverse complementarity on the intergenomic distances. And lastly, two artificial DNA sequences were generated by i) scrambling the T7 genome with Shuffle DNA, part of the Sequence Manipulation Suite (Stothard 2000) and ii) using Vladimír Čermák’s Random DNA Sequence Generator at http://www.molbiotools.com/randomsequencegenerator.html to generate a 39937 bp (48.4%GC) sequence.

## Results and discussion

VIRIDIC calculates intergenomic similarities between pairs of viral genomes based on BLASTN alignments. Because these alignments depend on the BLASTN parameters used, we have tested four sets of such parameters (Fig. 1). These ranged from “relaxed” (-word_size 7 - reward 2 -penalty −3 –gapopen 5 –gapextend 2) to “very stringent” (-word_size 28 -reward 1 - penalty −2). All four sets of parameters performed similarly for genomes with a higher degree of similarity. However, for more distant genomes, the calculated similarity values were significantly lower for the “very stringent” parameters, compared with the “relaxed” ones (see Fig. 1). This difference was expected, because at “very stringent” parameters BLASTN produces alignments only for highly similar genomic regions, and thus the regions of lower similarity are not taken into account when calculating the intergenomic similarity values.

**Figure 1:**
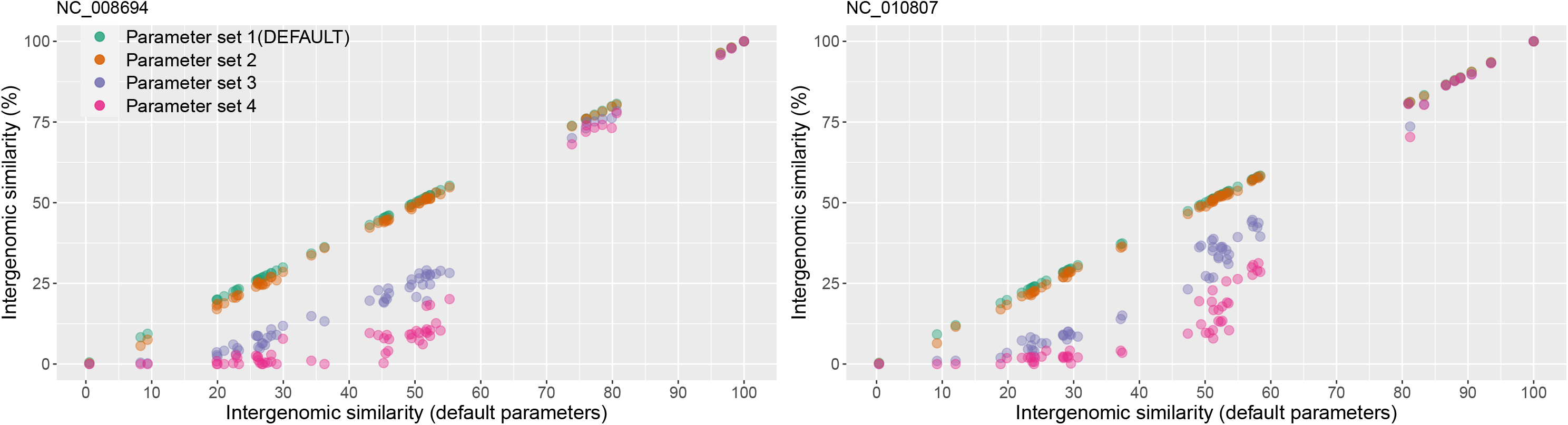
Comparison between the intergenomic similarity values produced with the default BLASTN alignment parameters (parameter set1: -word_size 7 -reward 2 -penalty −3 –gapopen 5 –gapextend 2) and parameter sets of increasing stringency. Parameter set 2: “-word_size 11 - reward 2 -penalty −3 –gapopen 5 –gapextend 2”. Parameter set3: “-word_size 20 -reward 1 - penalty −2”. Parameter set4: “-word_size 28 -reward 1 -penalty −2”. For illustration, the similarity values between two viral genomes (NCBI accession NC_008694 and NC_010807) and all the other genomes in the benchmarking dataset were chosen. On the X axis are plotted the intergenomic similarity values as calculated with the parameter set1. On the Y axis are plotted the intergenomic similarity values as calculated with each of the four parameter sets. The plot was generated with the ggplot2 R package (Hadley 2016).

Taking the above findings into consideration, we have chosen the “relaxed” parameter set 1 as the default for VIRIDIC. The other, more stringent parameter datasets are made available for the user because they significantly decrease computational times, which can be most advantageous when desiring to cluster at a high similarity thresholds (e.g. 95% for the species level) a large number of viral genomes, as for example found in viral metagenomic studies. On a benchmarking dataset of 61 phage genomes, VRIDIC needed 270 seconds with the default parameters, and only 56 seconds with the most stringent parameters. Because in the range 90%-100% intergenomic similarity, the “very stringent” parameters produced only a small decrease in similarity (see Figure 1), these parameters could be used in viral metagenomic datasets to enable clustering at species level (discussed also below).

Further, we have compared the intergenomic similarity values produced by VIRIDIC with those calculated manually from BLASTN alignments (the “traditional” method used by ICTV for phage classification) and those calculated by different other tools (see Figure 2). VIRIDIC showed the highest agreement with the manually calculated similarity values, being able to correctly indicate genome pairs with low similarity. In contrast, most of the other tools either gave artificially high similarity values for distant genomes (OrthoANI, SDT, EMBOSS Stretcher), or they significantly deviated from the traditional method (PASC, Gegenees BLASTN 5%), or they were not linear with respect to a type species (Gegenees BLASTN 0%). The artificially high similarity values were likely due to their calculation only for the aligned part of the genomes. When the intergenomic similarity is normalized only to the alignment length, even if only a small region is aligned between two genomes in a pair, the outputted similarity can be high. Instead, VIRIDIC normalizes the number of aligned bases between the two genomes in a pair to the lengths of both genomes, and thus estimates better the similarity between distant genomes. One such example is the pair between the genomes of Escherichia coli T7 phage and Pelagibacter phage HTVC011P. When visualizing the alignment between these two genomes (see Figure 3), it is clear that only a small portion of their genomes align. VIRIDIC reported a 0.34% similarity for this genome pair. However, other tools reported similarity values of 18.13% (PASC) and even > 42% (SDT and Emboss Stretcher), see SI Table 1. ANI tools have been extensively used in bacterial classification (Goris *et al.* 2007; Lee *et al.* 2016), and to a certain degree in phage classification (Accetto and Janež 2018; Oliveira *et al.* 2019), because their results mimic those of nucleic acid hybridizations. In our study, OrthoANI gave a similarity value of zero between the T7 phage and the Pelagibacter phage HTVC011P or the two artificially generated sequences. However, for the rest of the T7 phages the OrthoANI values plateaued at an artificially high 62% similarity, in agreement with the previous observation that ANI values below 75% are meaningless (Rodriguez-R and Konstantidinis 2014).

**Figure 2:**
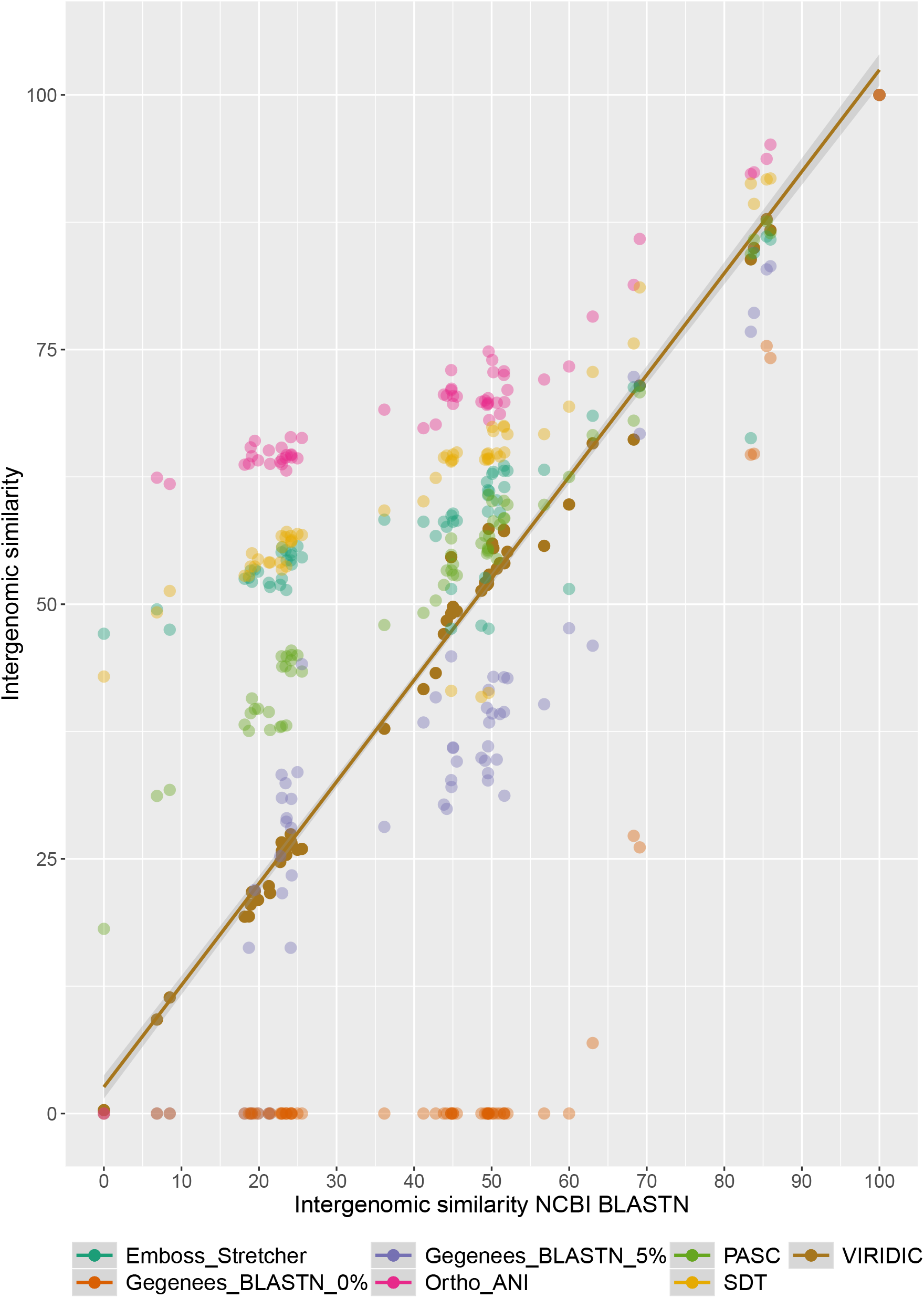
Plot comparing intergenomic similarity values generated by different tools (on the Y axis) with those generated by the traditional method used by ICTV (on the X axis). The plot was generated with the ggplot2 R package (Hadley 2016). Data used for this plot are found in SI Table 1.

**Figure 3:**
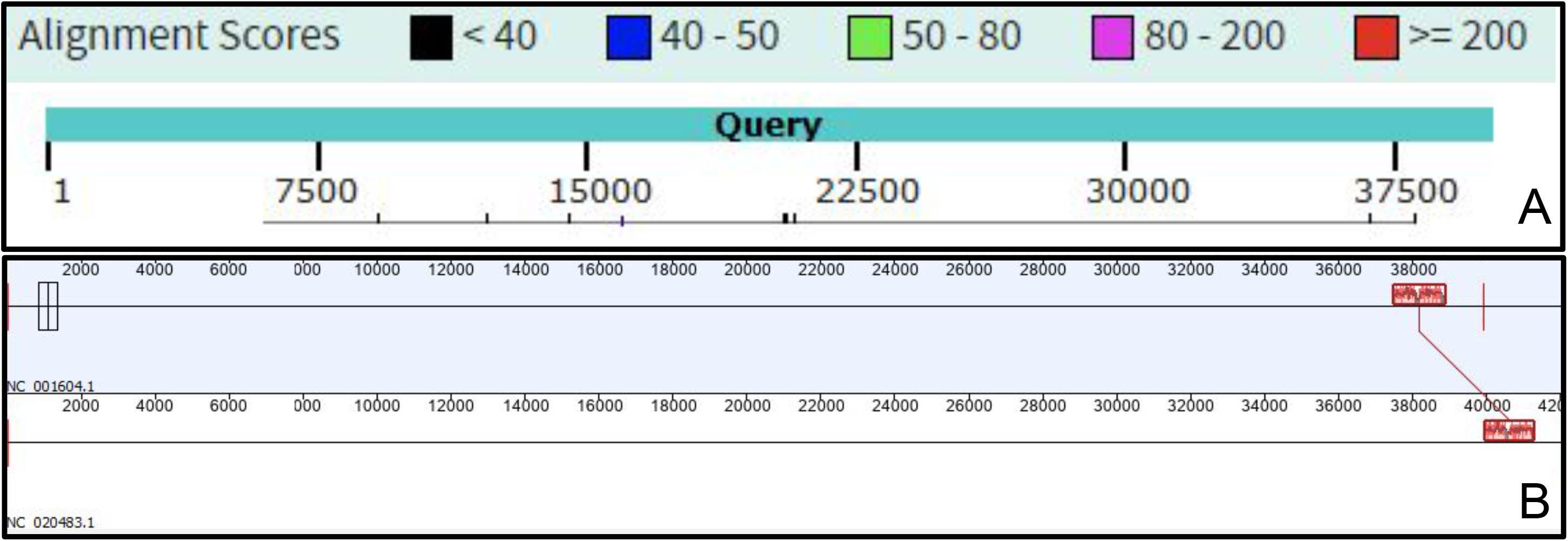
Genome alignments of the Escherichia coli T7 phage (NC_001604.1) and Pelagibacter phage HTVC011P (NC_020483.1) using A) NCBI BLASTN, with the T7 genome as query; and B) progressive MAUVE plugin from Geneious software (Kearse *et al.* 2012).

Following the calculation of the intergenomic similarities, VIRIDIC clusters and graphically represents these in a heatmap visualization (see Figure 4). Due to the color coding, groups of related phages can easily be recognized visually. Furthermore, if the results of the clustering are not satisfying, different clustering methods can be tested without recalculating the intergenomic similarities, which are the most time consuming. This is especially easy in the web-service version of VIRIDIC, which provides access through a graphical interface to many parameters for clustering and heatmap visualization. It is important to note that, although VIRIDIC performs a hierarchical clustering, the resulting tree is not a representation of the evolutionary paths and evolutionary distances between the different phages. To avoid such confusions, the tree resulted from clustering is only used to generate the heatmap and not visualized along its side. To reconstruct the phylogeny between or within the different virus clusters identified with VIRIDIC, further complementary phylogenetic analyses (e.g. core protein phylogeny) should be performed (Barylski *et al.* 2020).

**Figure 4:**
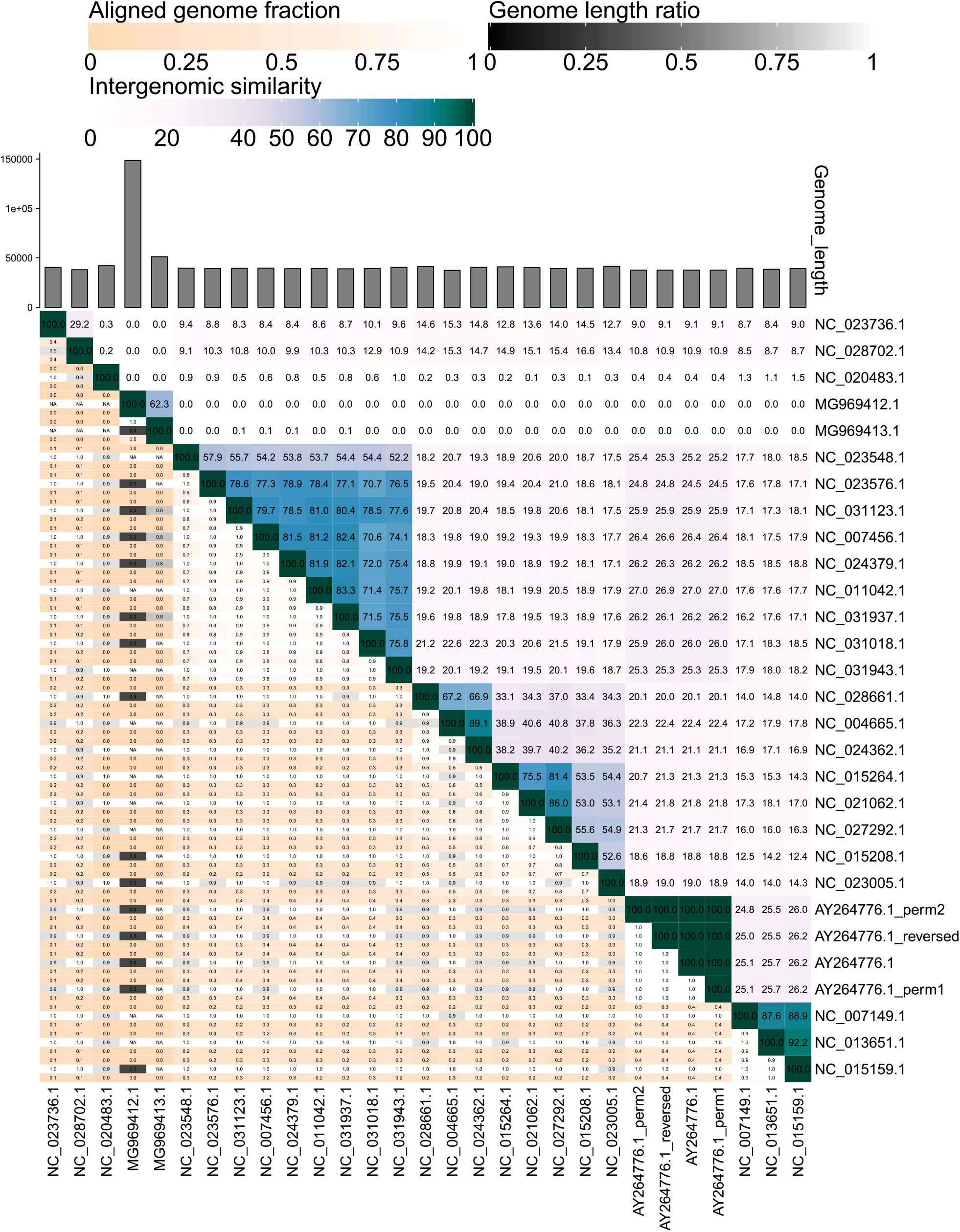
VIRIDIC generated heatmap incorporating intergenomic similarity values (right half) and alignment indicators (left half and top annotation). **In the right half**, the color coding allows a rapid visualization of the clustering of the phage genomes based on intergenomic similarity: the closer related the genomes, the darker the color. The numbers represent the similarity values for each genome pair, rounded to the first decimal. **In the left half**, three indicator values are represented for each genome pair, in the order from top to bottom: aligned fraction genome 1 (for the genome found on this row), genome length ratio (for the two genomes in this pair) and aligned fraction genome 2 (for the genome found on this column). The darker colors emphasize low values, indicating genome pairs where only a small fraction of the genome was aligned (orange to white color gradient), or where there is a high difference in the length of the two genomes (black to white color gradient). The aligned genome fractions are expected to decrease with increasing the distance between the phages. Therefore, darker colors should correspond to genome pairs with low similarity values, and whiter colors to genome pairs with higher similarity values. Similarly, closer related viruses are expected to have similar lengths. Therefore, if low genome length ratios correspond to genome pairs with high similarity (e.g. MG969412.1 and MG969413.1 have a 62.4% similarity, but only 0.3 genome length fraction), this signals that the pair needs to be investigated further before being classified. The genome of the K155 strain of the T7 phage (AY264776.1) and its permuted (AY264776.1_perm1 and AY264776.1_perm2) and reversed complemented (AY264776_reversed) variants presented no significant differences between their intergenomic similarity values.

In the heatmap, for display purposes, the similarity values have been rounded to the first decimal. This rounding can hide minute differences between almost identical phages. These differences will be visible however in the similarity table, another output of VIRIDIC, where the similarity values are represented up to the 3^rd^ decimal.

A third output of VIRIDIC is a cluster table, in which the phage genomes are grouped into putative species and genera, based on user set similarity thresholds (default 95% for species and 70% for genus). Because of difficulties associated with this threshold-based cluster detection, it can sometimes happen that the clusters in this file represent sub-clusters of species or genus level clusters easily identified by eye in the heatmap. This happens mostly to genomes with similarity values close to the species of genus threshold. Therefore, this table is not meant to be used instead of the heatmap in the process of phage classification. However, the cluster table can be useful when trying to de-replicate large datasets of viral genomes, in conjunction with the stand-alone VIRIDIC version.

When comparing viral genomes of different length, additional information can help to better interpret the intergenomic similarity values. This is the case especially when the two genomes in a pair share a high degree of similarity, as for example the pair between a complete and a partial genome, or between one smaller genome which is very similar to a region of a much bigger genome. For this purpose, VIRIDIC calculates three additional indicators of the alignment ability to capture the relatedness between viruses – the aligned fraction for genome 1, the length ratio between genome 1 and genome 2 and the aligned fraction for genome 2. Then it displays the three indicators in a color coded manner in the heatmap, as a visual aid for the user to spot genomes pairs of different lengths or partial alignments (see Figure 4). Even more, the length of each genome is plotted as an annotation along the columns of the heatmap.

The intergenomic similarity values calculated by VIRIDIC are not influenced either by genome permutations, or the genomes being in different directions (see Figure 4). However, the values will be influenced by the use of draft genomes at the scaffold level, which contain long stretches of “N”, because BLASTN is ignoring these regions. Therefore, it is not recommended to use such genomes.

The VIRIDIC web-service provides a graphical interface for running VIRIDIC remotely and it is meant for small to medium sized projects, ideally not bigger than 200-300 viral genomes. The stand-alone program can be run from the command line in Linux and thus, it can be integrated into bioinformatics pipelines. Furthermore, depending on the configuration of the computational resources, it can analyze a significantly larger number of viral genomes than the web-service. In the VIRIDIC workflow, there are two computational intensive steps, the BLASTN step and the calculation of intergenomic similarity matrix step. The computational requirements of the second step increase exponentially with the number of viral genomes to be compared. For example, we have run two projects, one with 169 and the other with 1236 viral genomes (size range 30 kb – 150 kb), on a Linux server with 40 central processing unit (CPU) cores and 256 GB RAM memory. The first project was finished in 10 min. The second project was finished in 19.5 h. For the BLASTN step, the number of CPU cores to be used can be controlled via a command line parameter. For the calculation of intergenomic similarity matrix, all available CPU cores will be used.

VIRIDIC offers several advantages compared to other similar tools. First, it provides a better estimation of the similarity between phages genomes, especially for the more distantly related ones. Second, it can be used in a high-throughput manner, allowing the analysis of datasets containing hundreds (the web-service) and even thousands (the stand-alone version) of phage genomes. Third, it generates an informative heatmap, which incorporates not only the similarity values, but also information about the genome lengths and aligned genome fraction, useful for evaluating the ability of the similarity values to capture the virus relatedness.

## Acknowledgments

We are thankful to Matthias Schroeder (Institute for Chemistry and Biology of the Marine Environment, Oldenburg) for excellent IT support through the project and for the preparation of the VIRIDIC singularity container.

## Table legends

SI Table 1. The complete list of viral genomes used for benchmarking VIRIDIC, their genome properties and intergenomic similarity values produced by different tools.

